# Cell-to-cell Modeling of the interface between Atrial and Sinoatrial Anisotropic Heterogeneous Nets II: Mechanisms Underlying Nets Dynamics

**DOI:** 10.1101/103259

**Authors:** Gabriel López, Norma P. Castellanos, Rafael Godínez

## Abstract

In [24] we studied the transitional zone between Sinoatrial cells and Atrial cells in the Heart. The present paper study the mechanisms underlying the dynamics of the modeled nets in that paper.

## 1. Introduction

Csepe et al. [7] study the functional-structural connection of the SAN and the atria. Their studies suggest that the microstructure of the connection paths between A and SA cells plays a crucial role in human SAN conduction and contributes to normal SAN pacemaking. Our paper may be considered as a local approach to the complexity of the specialized branching myofiber tracts comprising the SA connection paths described by Csepe et al. As an antecedent, we mention Benson et al. paper [3], where the authors study a finite, one-dimensional strand of cells with conduction patterns similar qualitatively to the model studied in the present paper. In their simulations a length step of Δ*x* = 0.2 mm was taken. This length corresponds to two cells of the size that we are considering, and consequently, a bigger approximation error in the numerical approximation, in comparison to the cell-to-cell model that we introduce in our example.

This paper is divided as follows: a) In section 5 we make a mathematical analysis of how it is possible to compare cell-to-cell models with partial differential equations (PDE) models. We reformulate the concept of liminal length in the context of cell-to-cell models, and establish that our approach is not opposite to Ruston’s liminar length concept for the cable equation, but in some sense complementary. b) In section 6 we state in Lemma 1, that the multidimensional parameter *α* = (*g*_12_,…, *g*_*ij*_,…, *g*_*nm*_) is a bifurcation parameter of a net of cells composed with *n* SA cells and *m* A cells coupled with conductance values *g*_*ij*_, *i* ≠ *j*, 1 ≤ *i* ≤ *n*, 1 ≤ *j* ≤ *m*. We show that the sum Σ_*ij*_ *g*_*ij*_ influences the behavior of the entire net in such a way that introducing conductance values as free parameters should lead to misleading conduction patterns in modeling nets. We give an elementary analytical proof of Lemma 1. We provide some arguments that prove that the geometry of a net is expressed implicitly by the non-zero conductance values and that they determine the change of qualitative behavior of the entire net. c) In 6.2 by using the piece-wise FitzHugh-Nagumo model we introduce elementary examples that illustrate the phenomenon stated in (a) and in (b).

## 2. General Materials and Methods

The equations used in [24] are of the form

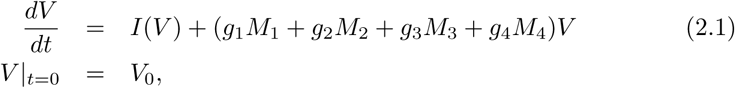

where the transposed vector *V*^*t*^ = (*V*_*A1*_,…, *V*_*An*_, *V*_*SA1*_,…, *V*_*SAm*_) corresponds to the Voltage (mV) of the A (*A*1 to *An*) and SA (*SA*1 to *SAm*) cells, with *n, m* values varying in different models, *I* (*V*)^*t*^ = (*I*(*V*_*A*1_),…, *I*(*V*_*SAm*_)), is a vector representing the currents (mV/ms) of A and SA cells, *M*1 is a connection matrix between A cells, *M*2 is a connection matrix between A and SA cells, *M*3 gives the connections between SA and A cells, and *M*_4_ is the connection matrix between SA cells; the constants *g*_1_ = *g*_*A−A*_/*C*_*mA*_, *g*_2_ = *g*_*A−SA*_/*C*_*mA*_, *g*_3_ = *g*_*SA−A*_/*C*_*mSA*_, *g*_4_ = *g*_*SA−SA*_. Here *CmSA* =.000032 *µ*F; *CmA* =.00005 *µ*F, and *g*_*SA−A*_, *g*_*A−SA*_, *g*_*A−A*_ are the conductance values between corresponding cells which values are specified in each model. The vector *V*|_*t*=0_ = *V*_0_, i. e. *V* at time *t* = 0, takes values which vary randomly with normal distributions, accordingly: for Atrial cells, mean −74.2525 mV, and for SA cells, mean −58mV, with standard variation .1 mV in both cells types. All the initial conditions in our models were taken from their respective papers, supplements, and codes provided by the authors when available. In tree models we keep *g*_*SA−A*_ = *g*_*A−SA*_ ≈ .6 nS, with the precise value given in the corresponding model.

The cell-to-cell approach requires: a) individual cell dynamics, modeled by Hodgking-Huxley type equations. For SA node cells, in [24] we used the model of Severi et al. [36], described in that paper A model of idealized two-dimensional arrangements of cells using a similar structure can be found in [39]; b) In order to implement better models, an approximate number of cells in SA node is required. This number may be estimated to be in the order of million, and we give an estimation in the Conclusions section.

Considering the cytoarchitecture of SA node not only the number of cells to which each cell is connected is important (already cited), but also the geometric distribution of each connection. Here we should remark that due to the use of connections matrices for the models of [24], the inherent three dimensionality of the cytoarchitecture (see [7] where the necessity of the 3D approach in order to understand the human SA node structure is amply discussed) does not require a special treatment as is the case for PDE, in which a tensor is required to describe the complex geometrical distribution of cells in the heart [11], [12], [13], [33].

Related to the integrating algorithms, we mention that stiff multi-step integration methods must be employed given the steepness of Atrial cells models. We used the quasi-constant step of Numerical Differentiation Formulas (NDF) method [21], in terms of backward differences, with absolute tolerance between two values *x, y* given by *|x − y|*=1*×*10^−12^, and relative tolerance: *|x − y|/* min(*|x|, |y|*)=1*×*10^−12^.

## 3. Results

Upon introducing numerical difference formulas in order to solve PDE we show that the resulting equations are equivalent to cell-to-cell ordinary differential equations systems with, of course, a corresponding approximation error. The simplest chessboard distribution geometry is the resultant pattern of such transformed PDE. So that those models are not interesting when studying the real cytoarchitecture structure of the SAN. With these transformed equations we can study the liminal length concept in cell-to-cell models. We obtained from this point of view that not only is a certain number of SA cells in a certain volume important when pacing the atria, but that the geometric distribution of the cells, also is. With this cell-to-cell view it is easy to see that our results are in some sense complementary to the Rushton’s liminal length. We are not interested in how big the minimum number of cells that activate a given region of the heart must be, but instead, how many A cells are activated by a given number of SA cells, and what their geometrical distribution should be in order to achieve conduction.

We found that the sum of conductance parameters and the geometric distribution of the conductance values of coupled cells through a net determine their change of qualitative behavior i.e., an entire net would be actively periodic or quiescent, according to the variation of such parameters.

## 4. Cell-to-cell vs PDE modeling

The simplest net models are formed with rectangular cells forming a chessboard-like geometric arrangement. In figure 1 the central cell is connected to two cells vertically, and two cells horizontally. The mathematical model of this net consists of a system of ordinary equations of the form

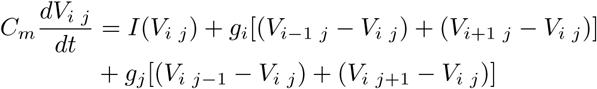

so that

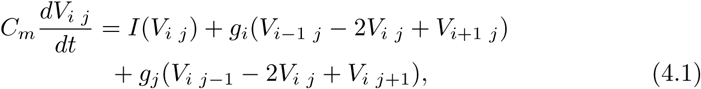

where *C*_*m*_ is the cell membrane conductance of each cell and *g*_*i*_, *g*_*j*_ are the coupling conductance, which, in order to simplify, we suppose to be different constants in the respective *i, j* subindex increment direction i. e., in vertical and horizontal directions respectively. Complexity may be added to the system allowing in the net different kinds of cells. For instance, in [45] two types of cells are included to form a mosaic model.

Let 1 ≤ *i* ≤ *N*, 1 ≤ *j* ≤ *M* and set *V*^*t*^ = (*V*_11_,…, *V*_1*M*_, *V*_21_,…, *V*_2*M*_,…, *V*_*N*1_,…, *V*_*NM*_) where *V*^*t*^ is the transpose of *V*. So, system (5.1) can be written in matrix form as

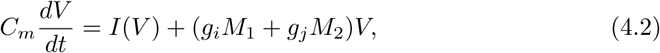

**Figure 1.**
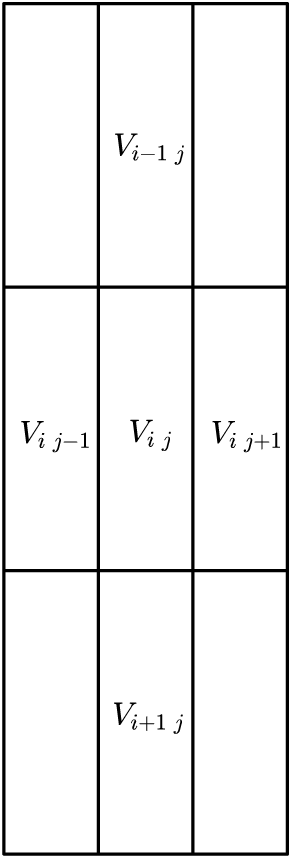
Scheme for simplest mosaic model. In this figure the central cell *V*_*ij*_ is connected to four other cells.

where *I*(*V*)^*t*^ = (*I*(*V*_11_),…, *I*(*V*_*NM*_)), *M*_1_ has the form

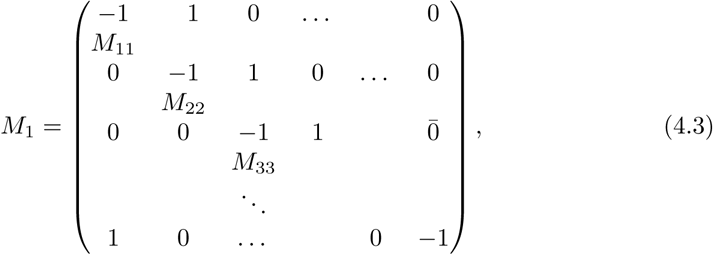

note that the rows containing ones correspond to the cells conforming the border of the arrangement, 0 are vectors of appropriate size containing only zeros, and *M*_*ii*_ are tridiagonal submatrices of the form

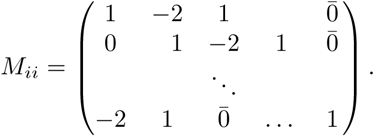

On the other hand, *M*_2_ is of the form

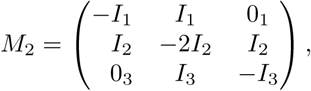

where 0_*i*_, *I*_*i*_ for *i* = 1, 2, 3 are zero and identity matrices of appropriate sizes. In order to introduce heterogeneity in the net by setting different types of cells distributed randomly, multiplying by a random matrix containing such information is required. Note that in this way, the resulting system is not equivalent to equations (2) and (3) in [45].

### 4.1. Mathematical Comparison between cell-to-cell and Macroscopic Models

From a mathematical point of view it is possible to convert a collection of ODE equations of a cell-to-cell model into a PDE corresponding to a macroscopic model, and reciprocally [21, sect. 4].To this aim to be achieved, first we recall the approximation formula for second derivatives 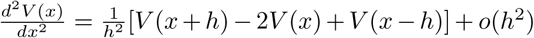 where *o*(*h*^2^) → 0 as *h* → 0. If we consider *h* = *l* of the order of cells length of course with the notation of the last section we have

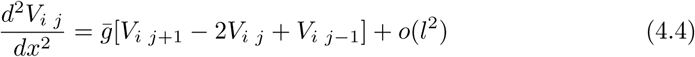

and

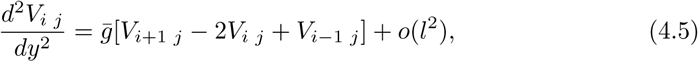

where *g* is a constant with appropriate units. Of course, the term *o*(*l*^2^) cannot be eliminated and hence *g*[V_*i*+1 *j*_ − 2*V*_*i* j_ + *V*_*i*−1 *j*_] would never be the second derivative of *V*_*i j*_ since *l* is supposed to be a constant and may be not reduced, in assuming that it is the minimum length of a cell. Given that the inaccuracy of equations (5.4) and (5.5) has been established, we avoid writing down the term *o*(*l*^2^) in the subsequent equations. Under these considerations we observe that mosaic model equation (5.1) is an approximation of the PDE

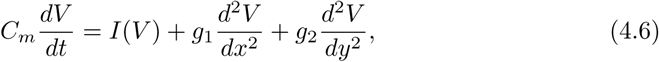

where *g*_1_, *g*_2_ are diffusion constants in appropriate units.

Reciprocally, we can associate with a PDE a system of equations corresponding to a certain cell-to-cell model. Actually, even the more accurate PDE’s models (after introducing numerical schemes) lead only to elementary cell-to-cell nets architecture and cannot predict properly complex phenomena such as slow conduction, decremental conduction, conduction block and changes in action potential duration and refractory periods, which are characteristics recognized as conditions playing an important role in causing reentrant arrhythmias [34]. Furthermore, those PDE approximations cannot model phenomena as the one described in [37] where the authors claim that myocardial architecture creates inhomogeneities of electrical load at the cellular level that cause cardiac propagation which is stochastic in nature.

### 4.2. Rushton’s liminal length

The concept of liminal length associated with the cable equation gives us an insight into what the minimum length of a segment with outward current in the finite border of a semi-infinite cable is, which prevents current to be inward. This may be interpreted as the size of a conductive tissue which may drive the heart as in [3]. Now we introduce the concept of liminal length in the cell-to-cell context. For instance, cable equation (16) in Noble’s paper [28] can be written as

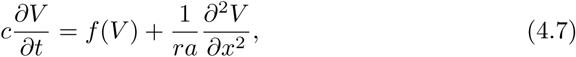

where 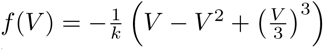. To introduce the cell-to-cell approach we use in (5.7) the approximation formula

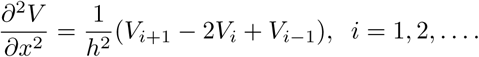

Let *V*_*i*_ the *i*-th cell in a series arrangement of an arbitrary number of cells. According to (5.7) each cell in the series satisfies

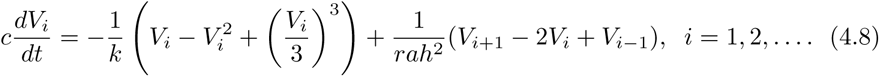

Set *g* = 1/(*rah*^2^). Now, for the entire series we may write the collection of equations (5.8) as a system of ordinary differential equations

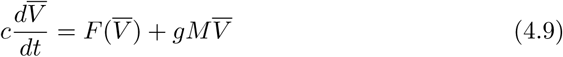

Where for a semi-infinite cable

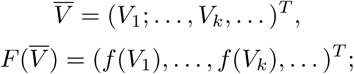

where *T* denotes the transpose of a matrix, and

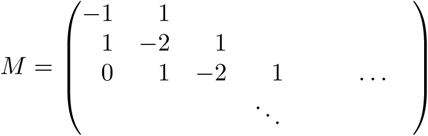

For a finite cable *i* = 1, 2*,…, k* we have then,

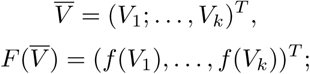

and now *M* is the finite matrix:

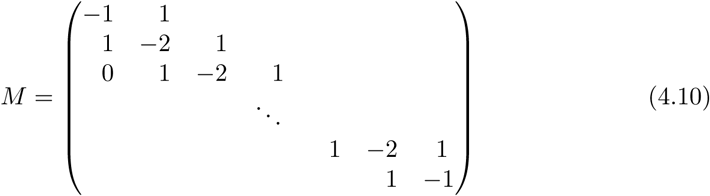

The most relevant point here is that system (5.9) corresponds to a *cell-to-cell* one variable per cell FitzHugh-Nagumo in a series system with cubic *f*. Hence, any analysis made in a semi-infinite cable equation (5.7). For instance the study in Noble’s paper will have an equivalent in the cell-to-cell model (5.9). As an example, formula (4, p. 575) in Noble’s paper for Rushton *liminal length x*_*LL*_:

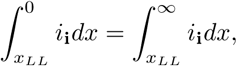

may be written as

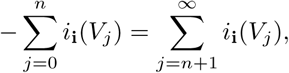

just taking *dx* = Δ*x* = *h*, where *h*, as before, is the length of each cell, and *n* is, hence, the minimal number of cells that provide inward current when the cable is at threshold.

#### 4.2.1. FitzHugh-Nagumo model

For the FitzHugh-Nagumo with *k* cells and with two variables in each cell we have the system

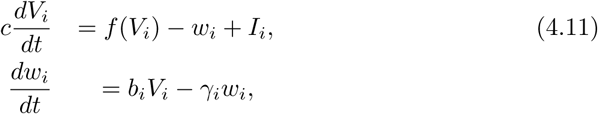

where *i* = 1,…, *k* and *f*(*v*) = *v*(*a* − *v*)(*v* − 1), 0 < *a* < 1, and 0 < *b*_*i*_, *γ*_*i*_ constants.This system can be written in a matrix form as

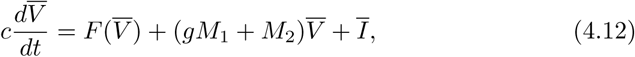

where *V, F, M*_1_, *M*_2_, whose exact form is described in the Simple example. In comparing equation (5.12) with (5.9) we note that the most notorious difference is the presence of matrix *M*_2_ which contains the bifurcation parameters of the FitzHugh-Nagumo cells model. In the section Discussion an extended analysis of some other differences is established.

With equations (5.7), (5.12) we can make an analysis of the liminal length concept included in Discussion in the context of cell-to-cell models.

## 5. Mechanism underlying Nets Dynamics

In this section, we state and prove Lemma 1 which claims that the sum Σ*g*_*ij*_ acts a bifurcation parameter of an entire system of cells, this is a first mechanism leading the net behavior. In the Simple example, we give a simple example that may clarify our arguments. A second mechanism is obtained by different geometric arrangements of cells that lead to a different characteristic polynomials of the net and, therefore to different set of bifurcations. A third mechanism for the change of behavior in the nets, focuses on bifurcation parameters of each cell, and how through the stimulation of neighboring cells, A cells switch from quiescent to active.

### 5.1. First and second mechanisms

The question we want to answer in this section is: why do different geometric configurations lead to a different net behavior?, why is it not enough to study series of cells, but three dimensional nets? In order to respond to these we introduce an abstract form of A and SA cells equations:

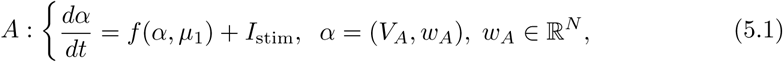

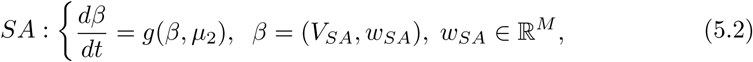

where *µ*_1_*, µ*_2_ are bifurcation parameters, possible multidimensional ones, each one, characteristic of the corresponding cell model; *I*_stim_ is a stimulus such that *I*_stim_ ≡ 0 leads to a quiescent state of A cells, note that in vitro, *I*_stim_ is usually constant or is given for a Dirac’s delta type function; and *V*_*A*_, *V*_*SA*_ ∈ ℝ are the voltage of corresponding cells. Generally, *N* ≠ *M* when modeling the transitional zone. Consider the characteristic polynomials

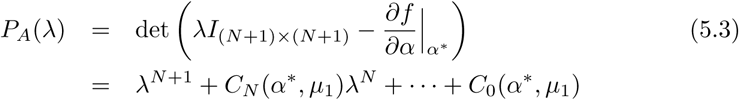

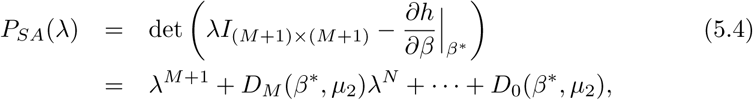

where *I*_*l×l*_ is the identity matrix of 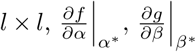 are the jacobians of *f* and *g* at equilibrium points *α*\*, *β*\* respectively. It is well-known that the change in qualitative behavior of the dynamical systems is given for the change in the roots of the characteristic polynomials as the coefficients *C*_*N*_, *C*_*N*−1_,…, *C*_0_, *D*_*M*_, *D*_*M*−1_,…, *D*_0_ change when *µ*_1_, *µ*_2_ respectively vary, for instance if complex roots of the polynomials change to purely imaginary ones, and so on.

In general, in coupling *k*, A cells with *l*, SA cells through _*g*_*ij*__, 1 ≤ *i* ≤ *k*(*N* + 1) + *l*(*M* + 1), 1 ≤ *j* ≤ *k*(*N* + 1) + *l*(*M* + 1) the characteristic polynomial of the system has the form

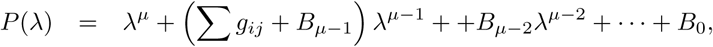

where *µ* = *k*(*N* + 1) + *l*(*M* + 1). In this case, the coefficients *B*_*j*_ depend on the coefficients of the characteristic polynomial of each cell and depend on each coupling *g*_*ij*_ as well. So that the sum Σ*g*_*ij*_ acts as a bifurcation parameter of the entire system and can be seen as a perturbation of a polynomial of degree *k*(*N* + 1) + *l*(*M* + 1) by a polynomial of lower degree. The affirmations of the last paragraph can be written as a lemma based on the following definition.

#### Definition

*We say that a system formed with n cells of the type A, SA in equations (6.1), (6.2) is conductance coupled if in each cell i of the system, the equation for* 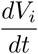 *includes the sum* Σ_*j*≠*i*_ *g*_*ij*_(*V*_*j*_ − *V*_*i*_) *for at least one j* ≠ *i*.

Note that in a conductance coupled system there are not isolated cells, i. e., each cell is connected to at least one other cell.

#### Lemma 1

*In a conductance coupled system formed with n cells of the type A, SA in equations (6.1)*, (*6.2*) *the sum* Σ_*ij*_ *g*_*ij*_ *is a bifurcation parameter of the system*.

The proof of the lemma is simple. Given that the system is conductance coupled, the term *tr*_*i*_ = Σ_*j,j*≠*i*_ *g*_*ij*_ appears in the mean diagonal of the jacobian matrix *J.* This is simply because in each equation for 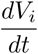, the following term does appear

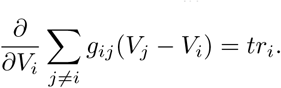

Therefore Σ_*ij*_ *g*_*ij*_ = Σ_*i*_ *tr*_*i*_ is part of the trace of *J*, and consequently, the characteristic polynomial of the system is of the form

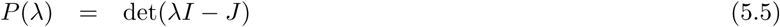

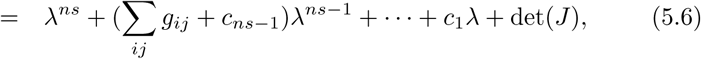

where “det” denotes the determinant of a matrix, and *s* is the sum of the number of variables of the SA and A cells in (6.1), (6.2).

Given that the trace of a matrix is invariant under the choice of basis and invariant under elementary transformations of matrices (as well as det(*J*)) the sum Σ_*ij*_ *g*_*ij*_ is a bifurcation parameter of the conductance coupled system, as we claim, because the coefficients (Σ_*ij*_ *g*_*ij*_ + *c*_*nk*−1_) and det(*J*) would determine the change of the number and characteristics of the real roots of *P* (λ).

#### 5.1.1. Second mechanism: the geometry of the net

In the general case, we can write the equation of the system in the form

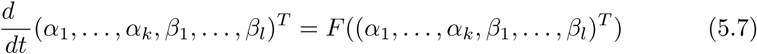

So that the vector containing all the cells may be relabeled as

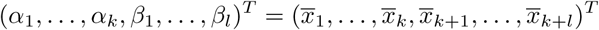

in such a way, the coupling *g*_*ij*_ ≠ 0 means that the cell *x*_*i*_ is coupled with the cell *x*_*j*_. With this the jacobian of the net has the form

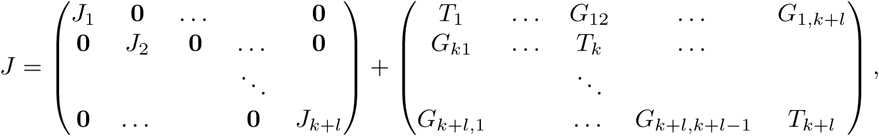

where *J*_*i*_ are the jacobian matrices of each cell *x*_*i*_, 0 is a zero block matrix, so that the matrix containing the *J*_*i*_, is block diagonal; *T*_*i*_ are block matrices, suited also as block diagonal, with *tr*_1_ in the first entry and zeros otherwise; and *G*_*ij*_ are block matrices with *g*_*ij*_ in the entry *ij* of the matrix and zero otherwise. The *T*_*i*_ blocks are formed as before by the derivative 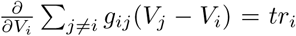. The blocks *G*_*ij*_ are formed by the derivative 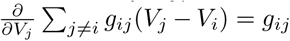, because of the definition of the jacobian of a system. Since the coefficients *c*_*r*_, *r* = 1,… *ns*, of polynomial (6.5) depend on *g*_*ij*_, so do the bifurcations, as simply varying *k* and *l* shows i. e. varying the number of SA and A cells. Furthermore, since det*J* may depend on *g*_*ij*_ (as well as all the other coefficients) the geometric distribution of cells depends on these parameters. Since the A and SA are cylindrical and the coupling is through cell membrane connexins, the net geometry is determined according to which parameters *g*_*ij*_ are different from zero.

Let’s call *J*_model_ the block diagonal matrix with diagonal blocks *J*_*i*_, and lets call *G*_geometry_ the matrix with *T*_*ri*_ and *G*_*ij*_ block so that we can write *J* as

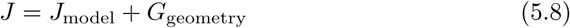

We conclude, regarding equation (6.8), that the second mechanism is given by *G*_geometry_, and the third mechanism that determines the nets dynamics is given by the bifurcation parameters of each cell in a net, given by matrix *J*_model_. This last mechanism is not less important than the others, but more difficult to detect. The so-called, global bifurcations, can be detected only by numerically integrating the equations of the systems (see for instance [16]). Hence, the bifurcation parameters treated in subsection 6.1 in many cases are less expensive in computational time than global bifurcations. We think that the more important bifurcations of complete nets in this paper are of this type. In any case, the way in which we found them, was by numerically integrating different geometric arrangements.

### 5.2. Third mechanism

In considering the jacobians of each cell in equation (6.8) we have to study the bifurcation set of each cell in a net at equilibrium points. In [16] just for the Hodgking-Huxley model (of four variables compared with thirty eight of LC model) four bifurcations of codimension one and six of codimension two are described. Thus, the number of bifurcations of codimension 1,2, et cetera in the LC model is too big to be treated computationally, not to mention analytically. With this in mind, we do not proceed further with such analysis. Given the set initial conditions for A cells (CI), which are determined experimentally, we are interested only in the bistability of such cells in a neighborhood of those CI. Consequently, we are interested only in the way in which A cells change from a stable quiescent state, to the active stable state through stimulation by neighboring cells. In considering LC model, for instance, if stimulus I is given as,

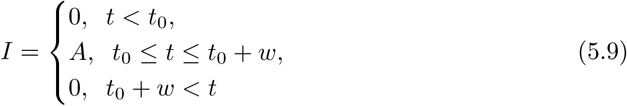

where *A* is the amplitude of the stimulus in picoAmperes, *t*_0_ = .1 s, is the starting time, and the duration of stimulus is *w* = 2 ms.

**Figure 2.**
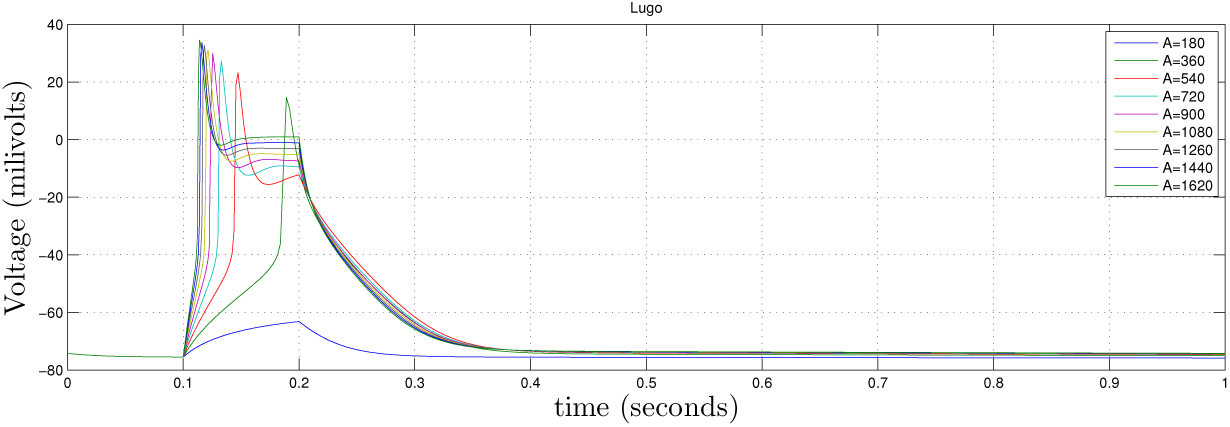
LC model threshold with I1. With amplitude of the stimulation of A=180 pA the cell is at subthreshold state, as mentioned in the text, the duration of the stimulus is *w* = 2 ms, but the activity in the cell (the little hump) lasts over 100ms. Note that with values at which the cell reaches threshold A ≥ 540 pA, the characteristic plateau of A cells is evident.

Then in Figure 2 we can see that LC model reaches threshold with the stimulus I taking *A* > 800 pA. In this way, an A cell would reach threshold if the sum of peaks of neighboring cells SA or A cell are about this quantity. Otherwise cells remain quiescent. When A cells model depolarizes they have a refractory period of about 300 ms as shown in the figure.

#### Simple Example

##### FitzHugh-Nagumo equations

Here we give the matrix form of a FitzHugh-Nagumo system used in section FitzHugh-Nagumo model. In connecting *in a series*, each cell in the system (5.11) with coupling conductance *g*, we obtain the system

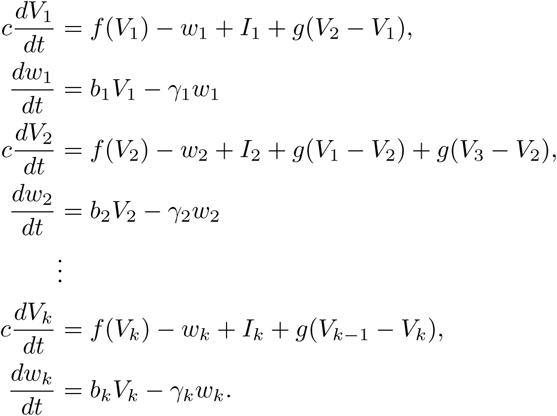

This system can be written in a matrix form as

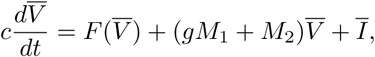

where

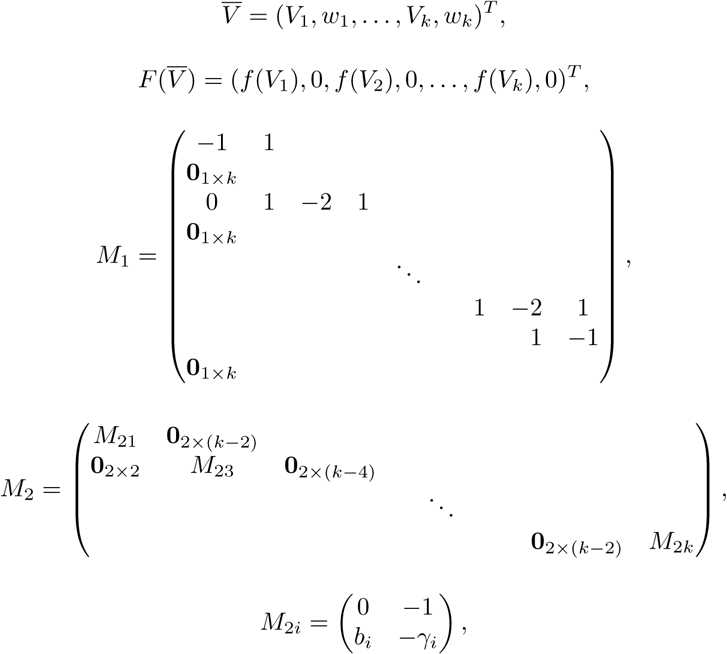

and *I* = (*I*_1_, 0, *I*_2_, 0,…, *I*_*k*_, 0)^*T*^. With equations (5.7) and (5.12) we make an analysis of the liminal length concept in the context of cell-to-cell models, and to compare the cable system equation with the FitzHugh-Nagumo model as in section Rushton’s liminal length.

##### A basic example

Now we give some estimations of how the sum of couplings *g*_*ij*_ may act as bifurcation parameter causing an entire net to collapse, even in the extreme case in which each cell in the net is periodic. Note that this case is extreme not only because we may think that if the coupling parameters are small, a given number of oscillators with identical dynamics could oscillate as an uncoupled system just because everyone would act as autonomic individual, but also, because the theory predicts that under certain conditions this kind of systems have an stationary state of the uncoupled case for *max*(*g*_*ij*_) small (see thm. 2.1 [26]).

To present the examples of this section we introduce two cell types, which caricaturize more complex cells models of SA and A cells. By using the piece-wise linear FitzHugh-Nagumo model it is possible to generate periodic autonomic cell models which we call SAFN and quiescent cells which we call AFN. For SAFN cells we use the equation (5.11) with *f*(*v*) = 1/6*v*, *I* = 19/72, *b*_*i*_ > 1/36 as an heterogeneity parameter and *γ*_*i*_ = 1/6. Now we summarize well-known facts of such a model: i) Due to Poincaré-Bendixon theorem SAFN cell posses an stable limit cycle, ii) due to Hopf bifurcation theorem the period *T*_*i*_ of each cell is approximately 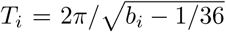, iii) the characteristic polynomial for these cells is *P* (*λ*) = *λ*^2^ + (*b*_*i*_ − 1/36), and hence it has two purely imaginary roots, iv) due to Torre theorem *n* cells do synchronize in phase for each *n* ≥ 2.

Now suppose we have *n* oscillators. To simplify the exposition we set *b*_1_ = *· · ·* = *b*_*n*_ = *b* and 0 < *α* = *b* − 1/36 and assume that each cell is connected with each other. Then the complete set of oscillators satisfies

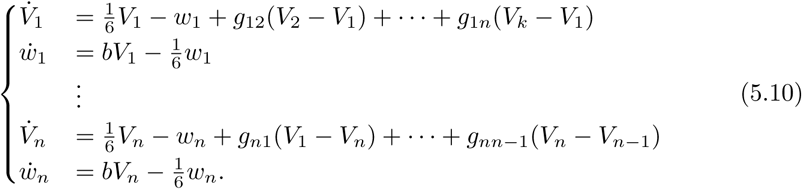

Denote Σ_*j,j*≠*i*_ *g*_*ij*_: = *tr*_*i*_, so the jacobian *J* of the system at (*V*_1_,…, *V*_*n*_) = 0 is

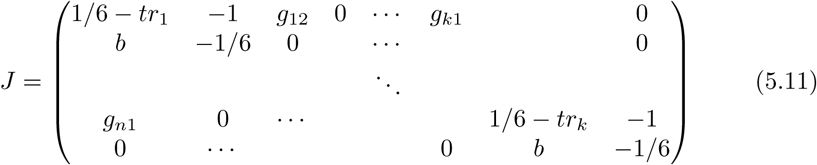

In this way the characteristic polynomial of the system is

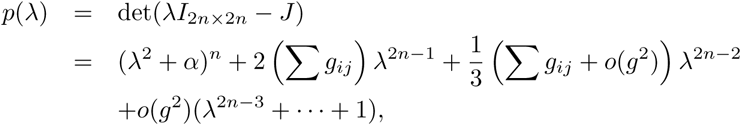

where *g* = *max*(*g*_*ij*_), *i, j* ≤ *n*. For weak coupling we may assume that *g* is small and therefore we can drop the terms of order *o*(*g*^2^). We write ϵ = Σ^*ij*^ *g*_*ij*_ and noticing that *ϵ* = *o*(*g*) we can see the characteristic polynomial of the system as an *ϵ*-perturbation with a polynomial of degree 2*n −* 1 of the polynomial (λ^2^ + α)^*n*^, and, actually, it can be shown that we can write

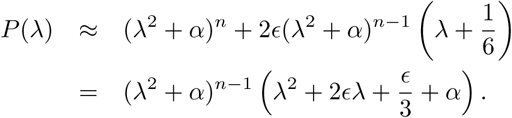

Now the polynomial (λ^2^ + α)^*n*^ has only complex roots meanwhile the polynomial with *ϵ* coefficient has one real root so for certain values of *ϵ* the polynomial *P* (*λ*) may change the number of real roots. Therefore, in that case the system would have a qualitative change of behavior, i.e. *ϵ* is a true bifurcation parameter of the system. In fact, 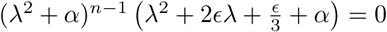 if and only if 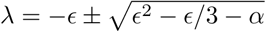 so that we would have real roots if 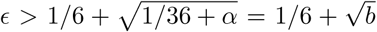. Therefore the entire net would have an stable fixed point if

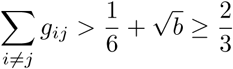

In modeling is usually taken *g*_*ij*_ = *g* as a constant for all *i, j*. In this case we obtain an estimation of the weak coupling which must be used with given *n*

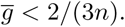

**Figure 3.**
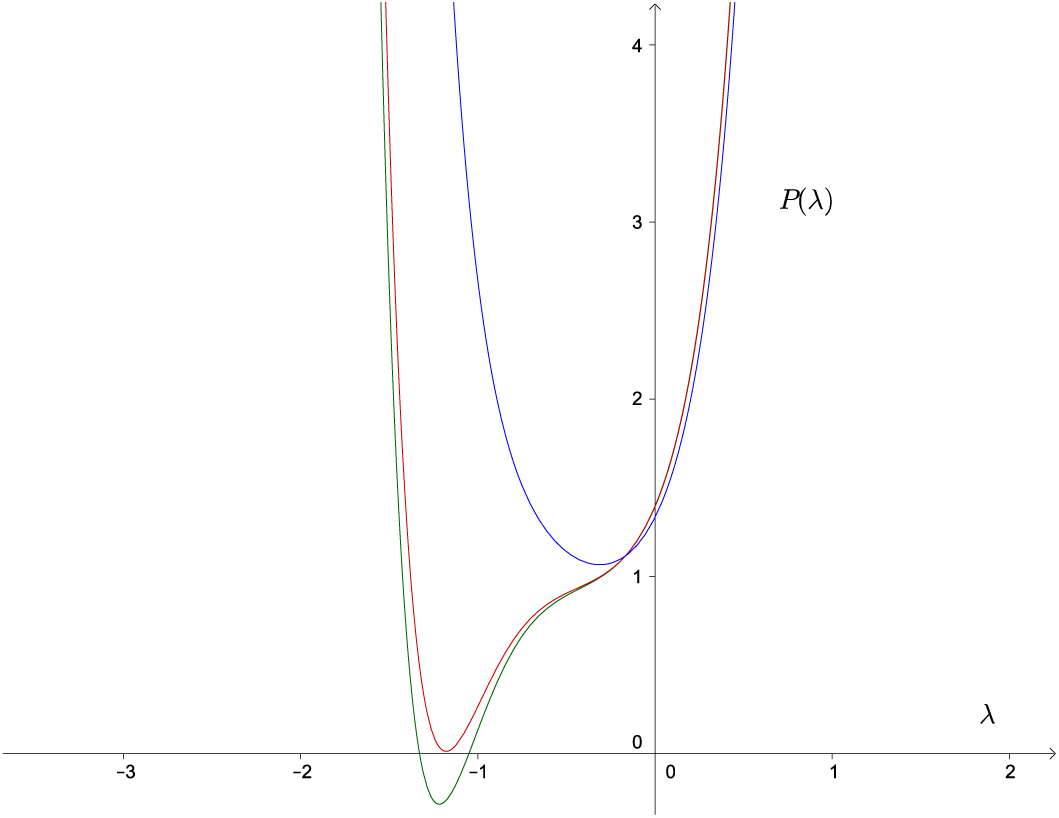
Perturbated polynomial *P* (*λ*). A polynomial of even degree with only complex roots is perturbed with a polynomial of lower order odd degree. The perturbed polynomial can have negative real roots and hence stable equilibrium points may appear trough bifurcation in the correspondent dynamical system.

More strikingly for a periodic net, if the *g* value is given (for instance experimentally) we obtain a bound for the number of cells in our net *n*, such that if *n* ≥ 2/(3*g*) the piece-wise linear FitzHugh-Nagumo model may collapse for certain set of initial conditions. Finally, we observe that in a more realistic model, many terms *g*_*ij*_ must be zero, otherwise each cell in the net would be connected with all and each other cells, which is not possible with cylindrical or spheric cells. But if one cell is connected with at least one other cell, their correspondent conductance will appear in the sum Σ*g*_*ij*_, and therefore may contribute in a possible collapse.

Even though the last model is very artificial it provides some nice insights. For instance, if the coefficients were variable then a massive heart attack could happen, let say, if all the values are at their maximum because, for instance, nervous system stimulation. On the other hand, introducing artificial big values of *g*_*ij*_ would make a failing net to conduct, for instance if and odd degree polynomial is perturbed by an even degree polynomial, introducing complex roots in the coupled polynomial which otherwise do not exist.

We show in the following example how the characteristic polynomial of three FitzHugh-Nagumo type cells, two of them periodic and one excitable change according with the geometry of the net. Each arrangement has a different characteristic polynomial and consequently a different qualitative behavior according with their respective bifurcation parameters variation. Now we introduce piece-wise linear FitzHugh-Nagumo model for A cells, which we call AFN model.

For AFN cells we use (5.11) with

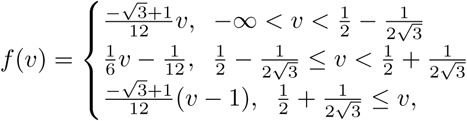

*b*_*i*_ = 1, *γ*_*i*_ = 1/6, *I*_*i*_ = *I*_*i*_(*ε*) = 1/12 + *ε*_*i*_. For this model it may be verified that if *ε*_*i*_ = 0 the cell would be quiescent. The reader may agree that systems with SAFN and AFN cells just mock any other physiologically accurate models. Nevertheless, such systems do capture some aspects that can be generalized to other more complex models. For instance the coupling of SAFN with AFN cells may change the quiescent of AFN cells to a periodic one acting qualitatively as a bifurcation parameter for AFN cells.

Lets denote the by *S*1 a series of two SA cells in which the AFN cell is connected only to one SAFN cell, by *S*2 we denote a series in which the AFN cell is connected to the two other cells. A ring of three cells is an arrangement in which each cell is connected to each other. These models are illustrated in figure 4. Note that in *S*1, *g*_*SA*2−*A*_ = *g*_*A*−*SA*2_ = 0 and the other conductance values are different from zero. For the ring all conductance values are not zero and for the series *S*2, *g*_*SA*2−*SA*1_ = *g*_*SA*1−*SA*2_ = 0.

Therefore any geometric arrangement in our example has a different characteristic polynomial of the form

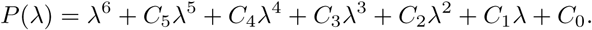

Coeffcients *C*_0_,…, *C*_5_ depend on the conductance values between cells and also depend on the parameters of each cell. In table 1 we compare the coefficients *C*_0_, *C*_4_, *C*_5_ of the three different groups in which *b*_*i*_ = 1 in (5.11). In every cell and all conductance values are equal to *g*, just to simplify the coefficients expressions.

It is possible to proceed further with these models, and to find values of *g* for which the characteristic polynomial of the net would have a pair of purely imaginary roots, (which is a necessary condition for having Hopf bifurcations) for instance applying Guckenhaimer et al. algoritms in [15]. In fact, theorem 2.5 in that paper gives necessary conditions which we can impose in *g* in order that the characteristic polynomial of a net to have exactly one pair of purely imaginary roots, which is a necessary condition in order to have Hopf bifurcations and hence, a probable periodic behavior in a net. Though, since this condition is necessary but not sufficient, after all the calculations in the mentioned theorem, in many cases in the net we cannot have the expected behavior.

**Figure 4.**
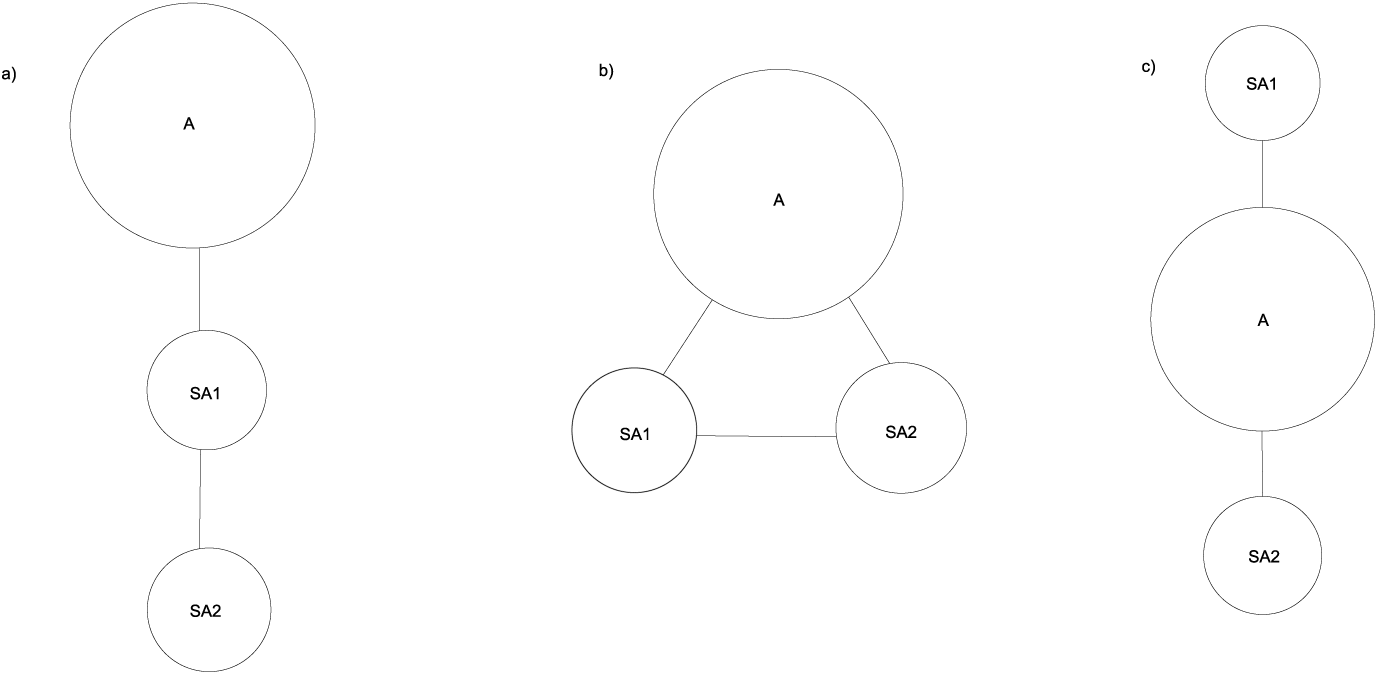
One AFN and two SAFN cells arrangements. In the figure a) is series *S*1, c) is *S*2, and b) is the ring. These are the simplest geometric arrangements with three cells with one A and two SA. The systems could have a very different dynamical behavior between each other according with given conductance values and according with the geometry of the net.

**Table 1.**
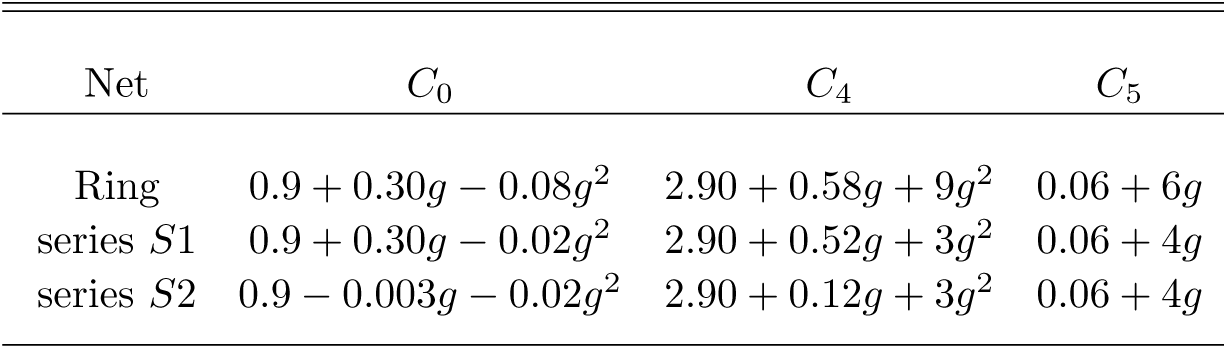
Comparing coefficients of characteristic polynomials

| Net | $C_0$ | $C_4$ | $C_5$ |
| --- | --- | --- | --- |
| Ring | $0.9 + 0.30g - 0.08g^2$ | $2.90 + 0.58g + 9g^2$ | $0.06 + 6g$ |
| series $S1$ | $0.9 + 0.30g - 0.02g^2$ | $2.90 + 0.52g + 3g^2$ | $0.06 + 4g$ |
| series $S2$ | $0.9 - 0.003g - 0.02g^2$ | $2.90 + 0.12g + 3g^2$ | $0.06 + 4g$ |

## 6. Discussion

### 6.1. Comparison of cable equation with FitzHugh-Nagumo model

When thinking that equation cable (5.9) represents the partial differential cable equation and FitzHugh-Nagumo equation (5.12) a minimal two variables cell-to-cell in a series system, we summarize some differences besides the obvious ones: i) Even that both equations can model a finite or infinite cable, we can add heterogeneity in (5.12) by simply choosing different *b*_*i*_, *γ*_*i*_. Moreover, this heterogeneity is dynamic; in fact, in choosing different parameters, it is possible to allow some cells to be active and some others to be quiescent. To say it in a picturesque way: cable (5.9) does not metabolize, whereas cable (5.12) at least captures, to some extent, periodic and quiescent behavior of A and SA cells. ii) Notice that the refractory period in cells may prevent outward or inward currents propagation, which cannot be modeled with the standard cable equation. Propagation may fail according to bifurcation parameters of individual cells, not only according to Rushton’s liminal length. For instance, *V*_1_(0) (the first cell *V* at *t* = 0) may be as big as anyone decides and current will propagate in (5.9), but for certain parameters *b*_*k*_, *γ*_*k*_, *k* > 1 within biological observations we may prevent propagation, regardless of the value of *V*_1_(0). So we may model a massive heart attack in this cable-heart with equation (5.12) but not with the standard cable equation. iii) We cannot use the steady state cable equation 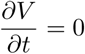, as *t* → ∞, since due to the periodicity of *V* in each cell, small scales of time are relevant in cell-to-cell modeling. iv) In comparing Benson et al. results [3] vs. our results, we remark the following: Benson et al. found that in a one-dimensional finite cable equation, conduction fails or succeeds according to: certain liminal length value, bifurcation parameters of the cells modeled with the paper’s equation, and certain change of transition parameters between cells. In order to compare the number of cells implicit in their numerical simulations from our standpoint, we note that the length step that they used (.2 mm) corresponds to one cell (which is not the observed size of SA or A cells, their approach is numerical not cell-to-cell) so that the 15 mm of one dimensional strand that they consider (p. 1320), corresponds to 75 cells in the cell to cell model. Authors used Lou & Rudy model with blocked I_NaK_ to simulate SA cells. Among other results, they conclude that the liminal length reduces from 17 cells to 15 cells (p. 1321 3.4 to 3 mm) with a linear change of parameters when the transitional zone es increased from 0 to 15 cells (0 to 3.0 mm). Some of their results coincide *qualitatively* with our analysis in one dimension, in the sense that they found that propagation may fail according to the bifurcation parameters of the cells. In other aspects, our analysis differs from theirs, given that we do not consider as necessary any transitional parameters between different kinds of cells. Moreover, we conclude that are a number of SA cells necessary to achieve conduction, but also, that certain geometrical arrangements are too, so that their 1D approach is not enough to obtain proper simulations.

### 6.2. Cell-to-cell vs PDE

Comparing modeling cell-to-cell vs. PDE we proved that introducing difference formulas in order to approximate PDE leads to very simple chessboard geometrical structures of ordinary differential equations (ODE) with only one variable, and sometimes with an approximation error bigger than individual cells’ dimensions. One advantage of using the equivalence of ODE systems with PDE, is that it does facilitate the interpretation of Rushton’s liminal length.Our approach is in a sense a complementary of Rushton’s liminal length; we are not interested in how many SA cells are required to activate a volume of A cells, but in how many A cells are activated by a certain arrangement of a small number of SA cells. Given the complexity of these systems, our analytical tools are not sufficiently powerful to determine this number. Analytic approach becomes very complicated even though the decomposition of the jacobian matrix *J* in (6.8) is very easy to obtain. From the dynamical systems point of view, this multiparametric matrix is very complex. Note that the matrix with *J*_model_ blocks depends on the cell models chosen, but *G*_geometry_ gives the relative position between cells independently of the models. So that if real structure of transitional zone were known Matrix *G*_geometry_ would remain fixed when certain accuracy of individual cell models would be achieved. We are not aware if nowadays this is the actual state-of-the-art in cell modeling. Accordingly, one may question how precise our geometric cells distribution is and how it depends on the A, SA models chosen. In deed, the characteristic polynomial of a system depends strongly on the models chosen and the exact one-to-one cell correspondence of our models compared with nature’s real heart could seem to be trivial. Although not as trivial as chessboard geometries included implicitly when difference derivatives formulas are introduced to solve PDE, and used even in an overwhelming number of papers using cell-to-cell modeling. For instance, we recall that the observed number of cells interconnected is 9.1*±*2.2 cells [17],but this number cannot be achieved in planar chessboard geometries extensively used in 1D and 2D models, nor in three dimensional blocks with a maximum of 6 cell connected to each other. More observational and experimental information is required on the specific transition zone in order to establish a correspondence between a big number of modeled cells and heart cells. Moreover, if the exact geometry of the SAN boundary would be known, it would provide us a criterion to compare models and to decide if they truly reflect observations, according whether their local nets nets collapse or not.

### 6.3. Local vs global approach

Another point that can be under discussion is how a local approach in the geometry may contribute to real modeling when considering that the sum of all the net conductances determines the behavior of the net. Here we think that groups of local nets with *appropriate behavior* can be considered as a computational syncytium (i. e. syncytium at least from a modeling stand point) and then, we may proceed by uniting different syncytium to form a mega-net, and so on. Upon regarding this question we have to estimate the complete number of cells in the atrium. The Orthof’s (1998) review [32] the author reports data sizes of SA nodes in some mammalians including humans. In there were reported humans’ SA node lengths of 15mm,7.3mm, and 9mm and corresponding widths of 5mm, 1.6mm, and 5 mm. More recently and with more refined techniques, in Chandler et al. [8] authors report 29.5 mm in length, 18 mm in height, and 6.4 mm in width which is the largest extension of SA node published, to the best of our knowledge. In Boyett et al. [4] for the rabbit, the authors estimate that there are approximately 5000 cells in 0.1mm^2^ of the center of SA node. With similar arguments and assuming roughly that the SA node and cells within the node are boxes, we can calculate the approximate number of cells in the entire human SA node with the data in [8], i. e. we obtain a SA node volume of 339.84 mm^3^, and assuming that 90% of this volume is connective tissue (as in the cat, see references within Boyett et al.), we get 33.98 mm^3^ of approximate total cell volume in the SA node. On the other hand, assuming that each cell has a volume of approximately 1000 *µ*m^3^ (37*±*3 and 5*±*0.2 *µ*m in length and width [10] we obtain a rough estimation of the number of cells in SA node is 34 million. Even if we consider only the so-called center of the SA node which is estimated as only 1% of the total of the SA node area [4], we obtain a gigantic number of calculations that surpass the computational capacity available technology today. This fact explains why the macroscopic PDE approach is still today the subject of major study. Apart from the number of cells, we need to improve our knowledge of the fine structure of the transitional SAN zone. In Chandler et al. [8] fantastic DTMRI/microCT data of the SAN are available, but the scales that they study are quite large to allow us to see the interdigitations in the transitional zone. Histology observations are known, for instance in Shimada et al. [37], and Hoyt et al. [17], which provide fantastic information of interconnections between cells, but not enough to make a one-to-one cell correspondence between a real heart and net models of big number of cells’ structures.

Another question which we can formulate is: how valid are models that consider only two types of cells? An improvement would be, as a first step, to introduce fibroblast dynamics [22] and their contribution within SAN and transitional zone geometry. After this, other observed cells types dynamics, for instance in [41] may be included. This is part of a future project which also may include nervous system stimuli simulations.

## 7. Conclusions

In order to extend the concept of liminal Length to 3D to the cell-to-cell approach, we found that not only are a number of cells necessary for conduction, but that the geometric arrangement is also relevant. Not only must the concept of liminal length be extended to, say, a liminal volume or a number of cells in a volume, but this concept must include an appropriate geometric distribution of SA and A cells.

In the literature some models of the transitional zone between SA and A cells require non-experimental conductance values in order to obtain propagation of the AP generated in the SA node. On the contrary, in [24] we obtained models which do not require artificial parameter values to achieve propagation. Moreover, according to our results, oversimplified 1D, 2D, 3D arrangements of cells, would make it possible to obtain conducting patterns that do not occur in real world by introducing conductance values as free parameters.

Our models mimic locally the strand structure (interdigitations) observed and studied in classical papers such as [44], and in recent papers such as [7]. Mathematically, our successfully conducting AP models may be explained by a change of qualitative behavior due to the change in bifurcation parameters associated with a correct source-sink relationship in the transitional zone. Remarkably, the bifurcation parameters give a geometrical distribution between A and SA cells which is compatible with interdigitating bundles structure of the transitional zone. We conclude that in modeling the cytoarchitecture of the transitional zone between SAN and Atrial zone it is essential for smooth propagation of AP to include geometrical arrangements, accordingly with some bifurcation parameters studied by dynamical systems theory.

